# Egocentric Perception of Walking Environments using an Interactive Vision-Language System

**DOI:** 10.1101/2024.12.05.627038

**Authors:** Haining Tan, Alex Mihailidis, Brokoslaw Laschowski

## Abstract

Large language models can provide a more detailed contextual understanding of a scene beyond what computer vision alone can provide, which have implications for robotics and embodied intelligence. In this study, we developed a novel multimodal vision-language system for egocentric visual perception, with an initial focus on real-world walking environments. We trained a number of state-of-the-art transformer-based vision-language models that use causal language modelling on our custom dataset of 43,055 image-text pairs for few-shot image captioning. We then designed a new speech synthesis model and a user interface to convert the generated image captions into speech for audio feedback to users. Our system also uniquely allows for feedforward user prompts to personalize the generated image captions. Our system is able to generate detailed captions with an average length of 10 words while achieving a high ROUGE-L score of 43.9% and a low word error rate of 28.1% with an end-to-end processing time of 2.2 seconds. Overall, our new multimodal vision-language system can generate accurate and detailed descriptions of natural scenes, which can be further augmented by user prompts. This innovative feature allows our image captions to be personalized to the individual and immediate needs and preferences of the user, thus optimizing the closed-loop interactions between the human and generative AI models for understanding and navigating of real-world environments.

## I. Introduction

Vision plays a critical role in human locomotion by providing feedback about the environment, allowing individuals to navigate obstacles, maintain balance, and adapt their behavior accordingly. Visual information helps in the coordination of motor actions, such as stepping over obstacles or adjusting walking speed [1], by allowing the brain to predict and respond to changes in the terrain. Without vision, humans often struggle to move efficiently and safely in dynamic environments, highlighting the fundamental link between visual perception and movement. Inspired by this biological vision-locomotor control, computer vision systems, powered by deep learning, have likewise been developed for robotics [2]-[7]. However, language can provide a more detailed contextual understanding of scenes beyond what computer vision alone can provide.

The use of image captions for human-robot locomotion is a relatively unexplored area of research. By translating complex visual information into natural language, image captions could provide a high-level abstraction of the real-world (e.g., terrain types and obstacles), in addition to providing an initiative user interface for human interaction. Textual information could be used by reinforcement learning algorithms to dynamically adapt the human and robot’s walking strategy, allowing for real-time optimization based on the described scene [8]-[10]. Recent advances in generative AI and vision-language models [11] have opened new opportunities for this area of research. For example, [12] and [13] recently showed that vision-language models could interpret dynamic and unpredictable environments and execute complex navigational tasks from natural language instructions for robotics.

Building on this recent progress, here we developed a new multimodal vision-language system for egocentric visual perception of real-world walking environments. We trained a number of transformer-based vision-language models [14]-[16], characterized by strong few-shot performance, which were pretrained on large vision datasets with natural language supervision [17]. Our system is able to generate accurate and detailed scene descriptions and identify obstacles, changes in terrain, and other critical navigational cues. We also developed a new speech synthesis model and a user interface to convert the generated text into audio feedback to users. Our system also uniquely allows for user prompts to tailor the generated image captions to the individual needs and preferences of the human user. To our knowledge, this is the first study to explore closed-loop human-AI interactions with vision-language models for real-world locomotion.

## II. Methods

### A. Image Dataset with Natural Language

We used the Microsoft COCO Captions dataset to finetune our vision-language models, which is a widely used benchmark dataset in computer vision and natural language processing [18]. The dataset contains over 200,000 images with human-generated captions. The images were collected from a wide range of contexts, including diverse settings such as urban environments and natural landscapes. For image captioning of walking environments, we adapted and manually curated a subset of the MSCOCO Captions dataset. We collected 43,055 image-text pairs based on the appearance of keywords. The average length of the captions in our new dataset was 10-15 words per caption. Prior to model training, we standardized the images to RGB format and performed a random split, allocating 90% and 10% of the images for training and testing, respectively.

### B. Vision-Language Models

Our image captioning module is used to generate detailed descriptions of the images using large vision-language models. We used an encoder-decoder framework (Fig. 1), which is common in state-of-the-art vision-language models, to translate images into text. The system extracts features from the initial input image and converts the 2D data structure into a sequence of visual tokens through a vision encoder. We then perform a sequence-to-sequence translation using our text decoder to produce contextually relevant captions corresponding to the visual environment.

**Fig. 1.**
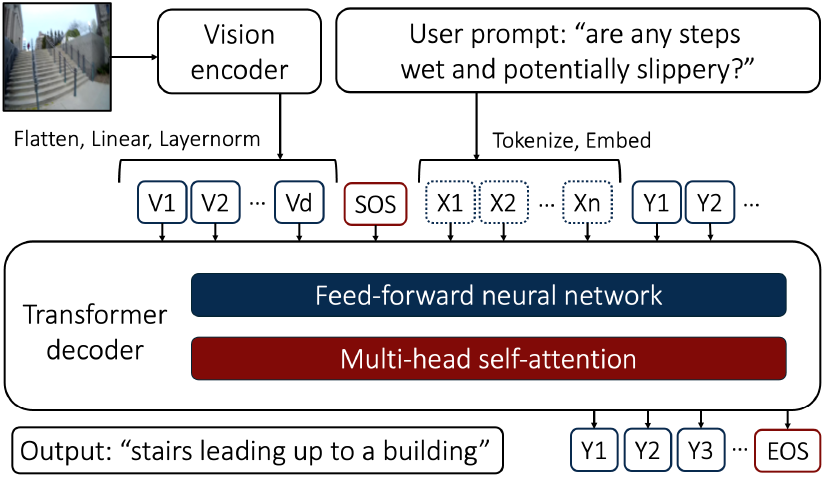
Our transformer-based encoder-decoder network architecture used for few-shot image captioning of real-world walking environments, including optional user prompts to tailor the generated image captions to the individual needs and preferences of the user.

This encoder-decoder framework was first proposed using a convolutional neural network as the vision encoder and a long short-term memory (LSTM) network as the text decoder [19] for real-world multimodal perception. To improve the quality of the text generation, we applied a transformer-based architecture to perform causal language modeling [20]. Causal language models can capture extensive context because they can attend directly to any previous token to predict the next token through the attention mechanism. They have consistently demonstrated impressive capabilities since the inception of large language models like GPT2 [15], which can capture extensive context because they consider all previous tokens to predict the next token. The transformer architecture is designed for parallel processing of sequences and uses attention mechanism to capture relationships among tokens, leading to a more coherent and contextually relevant text generation. Given a sequence of tokens, the general loss of causal language models is computed by:

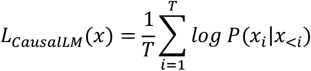

We first encoded the visual information into a sequence of embeddings analogous to word embeddings in language models. We used the Vision Transformer (ViT) model, which segments the image into patches for encoding [14]. Following the visual tokens, our transformer decoder starts the image caption generation with a start-of-sentence (SOS) token. To handle the multimodal nature of our task, we used bifurcated multi-head self-attention to separately model the interactions between the word and visual embeddings and the relationships among word embeddings themselves. This dual aspect of self-attention allows our models to produce captions that are both visually congruent and linguistically coherent. To model sequentially dependent structures, we masked the self-attention such that each caption token only attends to the preceding caption tokens and the visual tokens.

As shown in Fig. 1, our model also has the inherent flexibility to integrate user prompts as additional features for our image caption generation. If a user prompt is introduced, we initialize the decoding sequence after the visual embeddings and the SOS token, directing the decoder to generate the subsequent responses based on the user prompt. This novel function enhances the performance and human interaction for our multimodal system. We used two generative vision-language models as the backbone of our image captioning module, building on recent advances in vision-language pre-training [17], including the ViT-GPT2 and the Generative Image-to-text Transformer (GIT) models.

We first experimented with the ViT model as our vision encoder and the GPT2 as our text decoder [14]-[15]. The ViT model, designed to process visual data like sentences [14], splits the input image into a series of patches. These patches are then linearly embedded, with positional encodings added to retain the locational context, resulting in a sequence of visual embeddings that serve as an input to our language model. For text generation, the GPT2 model uses a robust transformer architecture to interpret these visual embeddings and generate descriptive captions. The pre-trained GPT2 comes with an extensive understanding of language and allows for relevant text generation that is synched with the visual input when adapted through fine-tuning on our new image-text pairs of real-world walking environments.

We also experimented with the GIT model [16] for image captioning. This unified transformer-based architecture uses the CLIP vision encoder based on contrastive learning between the vision and language modalities [17], which embeds visual tokens in the same embedding space as the language tokens, leading to a more consistent decoding process. The GIT model is pre-trained on a vast dataset comprising billions of vision-language pairs and demonstrates a strong capability to interpret complex visual environments and recognize scene texts without relying on external optical character recognition systems. We optimized the GIT model to provide descriptive navigational information.

We fine-tuned both vision-language models on our new custom dataset using an NVIDIA Tesla A100 GPU for 5 epochs with a learning rate of 5e-5. To reduce the high GPU RAM requirement due to the large model size and to accelerate model training, we used 16-bit floating precision for the numerical optimization on GPU instead of the commonly used 32-bit precision.

### C. Speech Synthesis Model

To convert the image captions from our vision-language models into speech for audio feedback to users, we developed a speech synthesis model building on state-of-the-art text-to-speech models. We used the SpeechT5 transformer encoder-decoder model as our backbone network with a set of modality-specific (speech/text) pre-nets and post-nets [21]. We used the text encoder pre-net to handle text inputs and the speech decoder pre-net and post-net for speech outputs. We used the text encoder pre-net to map the word tokens of our generated captions into a sequence of embeddings as inputs to our encoder. After the encoding, our speech decoder performs a sequence-to-sequence transformation to generate a log mel spectrogram conditioned on the hidden representations from the speech decoder pre-net and refined by the speech decoder post-net. We converted the output log mel spectrogram into an audio waveform via an additional vocoder neural network.

The SpeechT5 model can also convert into diverse voices, with the speaker embeddings as decoder inputs that capture a speaker’s unique voice characteristics [21]. We used X-Vector speaker embeddings, which are derived from deep neural networks and have shown to be superior in capturing contextual information from speech frames. Our speaker embeddings were sourced from the CMU ARCTIC speech synthesis database [22]. Our system provides users the ability to customize the output voice by substituting different speaker embeddings. The vocoder we used to generate the final audio waveform is an advanced generative adversarial network (GAN), which is designed to generate high-quality speech waveforms from spectrograms [23].

We designed a custom end-to-end pipeline to integrate our image captioning module with our speech synthesis module. The pipeline starts with our image captioning module, which takes visual input from the environment and extracts image features. Our vision encoder then transforms the features into visual tokens, followed by our text decoder to generate descriptive captions conditioned on an optional user prompt. This textual information is then rendered into speech waveforms by our speech synthesis model to provide users audio feedback of the system outputs, allowing the human user to interact bidirectionally with the generative AI models.

### D. User Interface Design

To enable human interaction and real-world practical use, we deployed our new multimodal system on the Gradio platform powered by cloud compute [24]. We configured the virtual environment required to support our models and all necessary scripts to execute the system. Additionally, we embedded our system in a web application based on the Next.js framework and hosted on GitHub Pages. We developed this web application with an emphasis on human interaction and a minimalist user interface design to promote accessibility across diverse user populations. Fig. 2 shows several examples of our user interface in action.

**Fig. 2.**
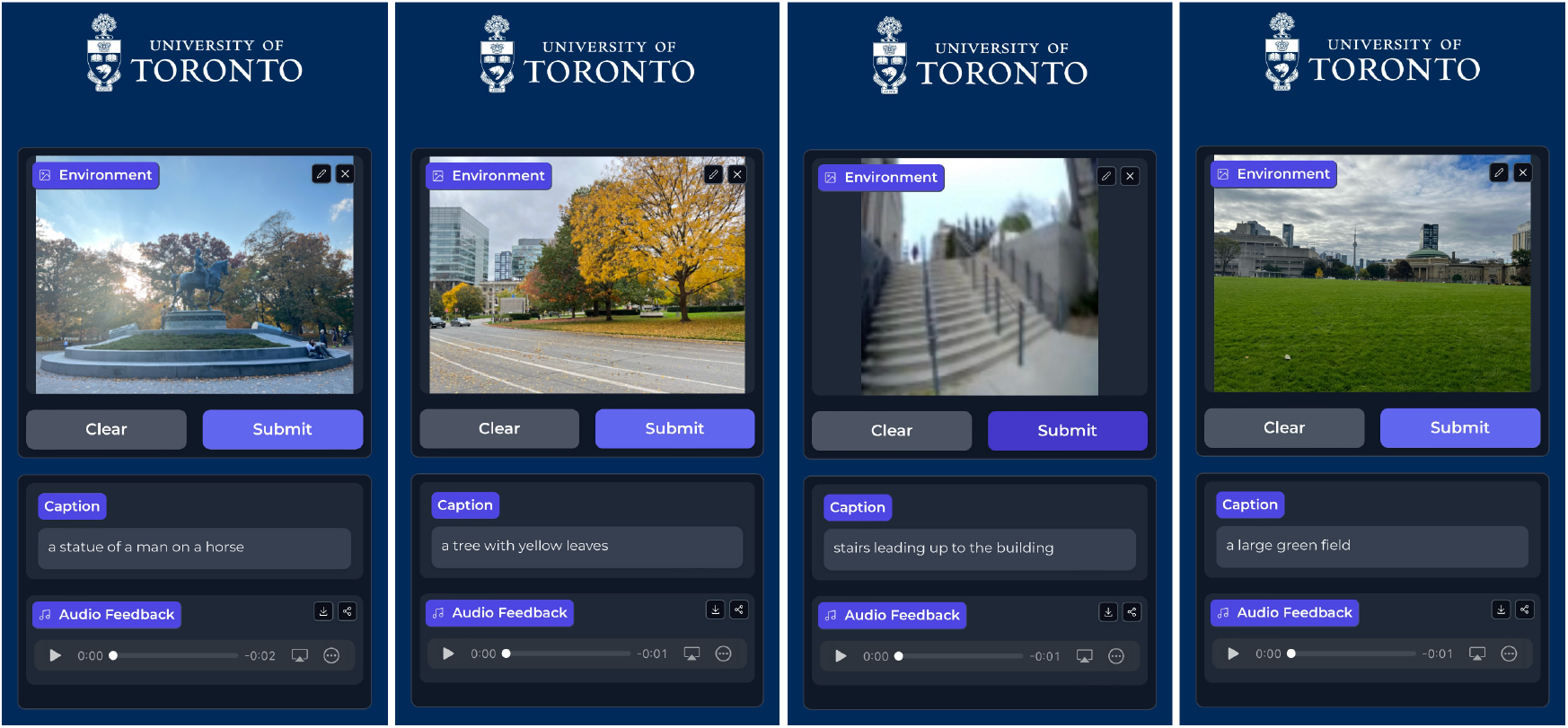
Our custom-designed user interface to provide audio feedback to users describing the generated image captions by the vision-language models. Our interface also supports feedforward user prompts, allowing for closed-loop human-AI interactions.

### E. Performance Evaluation

We quantitatively evaluated the performance of our vision-language models using several well-established metrics for text generation, including the Recall-Oriented Understudy for Gisting Evaluation (ROUGE) score. The ROUGE score measures the similarity between the generated text and a set of reference texts, which are typically human annotated. We focused on two common variants: ROUGE-N and ROUGE-L. The ROUGE-N score evaluates the overlap of n-grams, bordering sequences of *n* words, between the generated candidate and the reference text. We also used the ROUGE-L score that measures the longest common subsequence between the candidate and the reference. A higher ROUGE score indicates a better quality of the generated caption in terms of its similarity to the reference image captions that are human annotated. We also used the Word Error Rate (WER), which measures the number of substitution, deletion, and insertion errors in the generated image captions compared to the reference text. It is calculated based on 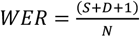, where *S* is the number of substitutions, *D* is the number of deletions, and *I* is the number of insertions needed to transform the generated image captions into the reference text. A lower WER score indicates better system performance.

We also tested the processing speed of our new system when deployed using cloud computing with different hardware for image captioning and feedback generation. We experimented with four different variants of our system: (1) Text-only feedback generation, (2) Text and audio feedback generation, (3) Text-only feedback generation with user prompts, and (4) Text and audio feedback generation with user prompts. We tested each of these four system variants with three different computing hardware configurations (Table 1).

**Table 1.** The different computing hardware that we used to test our new multimodal vision-language system for image caption generation.

| Computing | vCPUs | System RAM | GPU RAM |
| --- | --- | --- | --- |
| Intel Xeon CPU | 2 | 16GB | 0GB |
| Nvidia T4 GPU | 4 | 15GB | 16GB |
| Nvidia A10G GPU | 4 | 15GB | 24GB |

## III. Results

Our ViT-GPT2 vision-language model achieved a high ROUGE-1 score of 48.3%, a ROUGE-2 score of 21.8%, and a ROUGE-L score of 43.9% on the MSCOCO Captions dataset. Our GIT vision-language model achieved a low word error rate of 28.1%. In terms of deployed processing time (Table 2), our text-only feedback generation was the fastest across all computing devices, achieving speeds as fast as 2.2 seconds on the NVIDIA A10G GPU. Incorporating audio feedback led to slight increases in processing times (i.e., approximately 1 second). However, including user prompts significantly increased processing times, with average durations ranging from 11.3 to 15.3 seconds on GPU devices. The processing times varied depending on the computing hardware, with the NVIDIA A10G GPU consistently demonstrating faster processing speeds compared to the Intel Xeon CPU and the NVIDIA T4 GPU. Processing times on the CPUs were approximately 2-3 times longer than those on GPUs.

**Table 2.** The average processing times for our system variants, including (1) text-only feedback generation, (2) text and audio feedback generation, (3) text-only feedback generation with user prompts, and (4) text and audio feedback generation with user prompts. Each system variant was tested with three different computing hardware.

| System Variants | CPU | T4 GPU | A10G GPU |
| --- | --- | --- | --- |
| Text feedback | 7.5s | 2.4s | 2.2s |
| Text and audio feedback | 8.5s | 3.0s | 2.6s |
| Text feedback with user prompts | 30.2s | 12.5s | 11.3s |
| Text and audio feedback with user prompts | 31.1s | 15.3s | 12.4s |

## IV. DISCUSSION

In this study, we developed a novel multimodal vision-language system for egocentric visual perception, with an initial use case on understanding real-world walking environments. We trained a number of state-of-the-art transformer-based encoder-decoder models using 43,055 image-text pairs of scenes for few-shot image captioning. We also custom-designed a new speech synthesis model and a user interface to convert the generated text into speech for audio feedback to users. Most notably, our system uniquely allows for feedforward user prompts to personalize the generated image captions to the individual and immediate needs and preferences of the user, thus optimizing the closed-loop human-AI interactions for understanding and navigating real-world environments. Moving forward, this research has practical implications for robotics and embodied intelligence.

Our system is able to generate image captions with an average length of 10 words while achieving a high ROUGE-L score of 43.9% using the ViT-GPT2 model and a low word error rate of 28.1% using the GIT model. This is in comparison to previous image caption generation systems like [25], which score a bilingual evaluation understudy (BLEU) of 40.3% and a Metric for Evaluation of Translation with Explicit ORdering (METEOR) of 30.7%. One novelty of our system lies in finetuning the vision-language models on large-scale data specific to our application, which contrast previous research that simply applied pretrained models [13]. As mentioned by [26], fine-tuning can significantly improve the decision-making of vision-language models in tasks that require fine-grained visual recognition and visual semantic reasoning. These observations are in agreement with our study, which found that, after fine-tuning, the ROUGE score of the ViT-GPT2 model improved from 40.7% to 43.9%.

Unlike most control systems for robotics [27], our system uniquely includes the human in the loop by providing audio feedback as per the decision-making of the learning algorithm. While research in human-robot systems has included feedforward mechanisms for control [28]-[30], our system also provides feedback to the user. This integration can enhance user awareness compared to other systems, which could be beneficial for acceptance and trustworthiness of intelligent control systems. Our research increases the human interaction by including user prompts to tailor the generated image captions to the individual user needs and preferences. To our knowledge, this is the first study to develop and explore such capabilities for understanding and navigating real-world environments. As noted in [27] and [31], customizing robot behaviors to individual user preferences (e.g., while walking with a robotic prosthetic leg) is important for performance and our new multi-modal system provides a means by which the research community can explore such principles.

Despite these developments, our study is not without limitations. For example, one major challenge of our system is the real-time processing requirements of the large vision-language models, with increased computational costs associated with the user prompts and audio feedback generation. Including the user prompts significantly increased processing times, with average durations ranging from 11.3 to 15.3 seconds on GPU devices. This can limit the practical deployment of our system in resource-constrained environments where GPU computational resources are not readily available.

Moving forward, further research is warranted to improve our system efficiency. Optimizing the large vision-language models to reduce computational demands could help broaden the applicability and impact of our research. Exploring the potential for edge computing to handle the information processing locally could decrease the latency and reliance on cloud computing, thus making our system more practical in diverse applications. Moreover, adding an automatic speech recognition model [32] would be beneficial, where the closed-loop human-AI interactions could be performed through conversational feedback. In summary, here we developed a new multimodal, interactive vision-language system for egocentric visual perception, with an initial focus on understanding real-world walking environments. This system could be used to support research and development in robotics, embodied intelligence, and human-computer interaction.

## Acknowledgment

This research study is dedicated to the people of Ukraine in response to Russia’s ongoing invasion.

